# Dynamic recovery from depression enables rate encoding in inhibitory synapses

**DOI:** 10.1101/379081

**Authors:** Shiyong Huang, Morgan S. Bridi, Alfredo Kirkwood

**Author notes:** Correspondence (SH) (AK).

## Abstract

Fast-spiking parvalbumin positive interneurons (PV-INs) are essential for controlling network firing and the gain of the cortical response to sensory stimulation. Crucial for these functions, PV-INs can sustain high frequency firing with no accommodation. However, PV-INs also exhibit short-term depression (STD) during sustained activation, which is largely due to the depletion of synaptic resources (vesicles). In most synapses the rate of replenishment of depleted vesicles is constant, determining an inverse relationship between the STD level and the activation rate, which theoretically, severely limits rate coding capabilities. We examined STD of the PV-IN to pyramidal cell synapse in the mouse visual cortex, and found that in these synapses the recovery of depleted resources is not constant but increases linearly with the frequency of use. By combining modeling, dynamic clamp and optogenetics, we demonstrated that this dynamic regulation of recovery enables PV-INs to reduce pyramidal cell firing in a linear manner, which, theoretically, is crucial for controlling the gain of cortical visual responses.

## INTRODUCTION

In cortex the main mechanism that limits cortical activity is the inhibitory action of parvalbumin-expressing fast-spiking cells (PV-INs). Indeed, the recruitment of PV-INs reduces the spiking activity of cortical pyramidal cells in an additive and multiplicative fashion (Tremblay et al., 2016). Concordant with this function, PV-INs can fire at high rates, without attenuation, and are highly interconnected with pyramidal cells. However, like most cortical synapses, synapses made by PV-INs exhibit marked short-term depression (STD) of presynaptic release (Abbott et al., 1997; Zucker, 1989; Zucker and Regehr, 2002). Since STD strongly limits synaptic efficacy in the temporal domain, unraveling its rules and mechanisms is crucial for understanding neural processing by PV-INs.

Experimentally, STD is typically revealed as a progressive, yet reversible, depression of synaptic release that occurs during a train of repetitive stimulation. Conceptually, STD can be understood in terms of use-dependent depletion of synaptic resources. In the simplest models synaptic depression results from the depletion of the readily-releasable pool (RRP) of synaptic vesicles (Alabi and Tsien, 2012; Zucker and Regehr, 2002). Released vesicles are retrieved from the terminal surface followed by refilling with neurotransmitter via a process of ‘kiss-and-run’ (He et al., 2006) and/or clathrin-mediated endocytosis (Granseth and Lagnado, 2008). The steady-state amplitude of synaptic depression during a stimulation train reflects the balance between vesicle release and recovery.

In cortex, both excitatory and inhibitory synapses often exhibit STD, but with different properties and consequences. In most connections between pyramidal cells STD during repetitive stimulation follows the 1/f rule. That is, at the steady state the synaptic response amplitude decreases in inverse proportion to the presynaptic firing frequency (Abbott et al., 1997; Galarreta and Hestrin, 1998; Tsodyks and Markram, 1997). According to the depletion model, a 1/f rule could result when the vesicle release and recovery is constant and independent of presynaptic activity, as reported in hippocampal synapses (Wesseling and Lo, 2002). A cardinal consequence of this 1/f rule is that increases of the presynaptic firing rate above a certain limiting frequency do not result in commensurate increases in postsynaptic firing (Abbott et al., 1997; Tsodyks and Markram, 1997). This low pass filtering limits rate coding in these synapses, but makes them more suitable for temporal coding (Abbott et al., 1997; Grande and Spain, 2005; Tsodyks and Markram, 1997) and the detection of coincidences and synchrony (Cook et al., 2003; Reyes et al., 1996).

In contrast to excitatory synapses, inhibitory synapses, particularly those formed by parvalbumin-expressing fast-spiking interneurons (PV-INs), do not follow the 1/f rule. Instead, the relationship between the steady-state amplitude and the stimulation frequency is linear in inhibitory synapses in several brain regions, such as layer 5 of the rat cortex (Galarreta and Hestrin, 1998), striatal medium spiny neurons (Gittis et al., 2010), and layer 2/3 of rat visual cortex (Varela et al., 1999). The mechanisms underlying this linear relationship, and their functional consequences, remain largely unknown. By combining whole-cell recordings in connected PV-interneuron to pyramidal cell (PV→Pyr) pairs, dynamic clamp, and optogenetics, we revealed that use-dependent dynamic recovery from synaptic depression accounts for the linear relationship between frequency of presynaptic firing and postsynaptic responses. We also showed that this dynamic recovery enables a rate coding for the inhibitory function of PV-INs, that is, increasing the firing rate of PV-INs causes a linear reduction of firing in pyramidal cells.

## RESULTS

The 1/f rule, in which the amplitude of postsynaptic currents at the steady state is inversely proportionate to the frequency of presynaptic activity, is common among depressing excitatory synapses (Abbott et al., 1997), but not among depressing inhibitory synapses (Gabernet et al., 2005; Galarreta and Hestrin, 1998; Gittis et al., 2010; Varela et al., 1999). In a resource-depletion model of synaptic depression, the 1/f rule arises as a consequence of there being a constant rate for the replenishment of depleted resources. Thus a deviation from 1/f would suggest that recovery somehow depends on the frequency of presynaptic activation. We examined this possibility in synapses made by PV-INs onto layer 2/3 pyramidal cells in visual cortical slices.

### Different dynamics of EPSCs and IPSCs in layer 2/3 pyramidal cells of visual cortex

First we confirmed that previously-reported differences in the dynamics of excitatory postsynaptic currents (EPSCs) and inhibitory postsynaptic currents (IPSCs) found in the somatosensory cortex can also be found in the visual cortex (Figure 1A). During prolonged repetitive stimulation the IPSCs and EPSCs recorded in the same pyramidal cell both depress, but with different frequency dependencies (Figure 1B-C). The average depression level at the end of a long train roughly followed the 1/f rule in the case of EPSCs, whereas the relationship with stimulation frequency was better fitted with a linear function in the case of IPSCs (Line: R^2^ = 0.996, 1/f: R^2^ = 0.164, regression; F(1,3) = 670.3, p = 0.0001, Extra sum-of-squares F test; Figure 1D).

**Figure 1.**
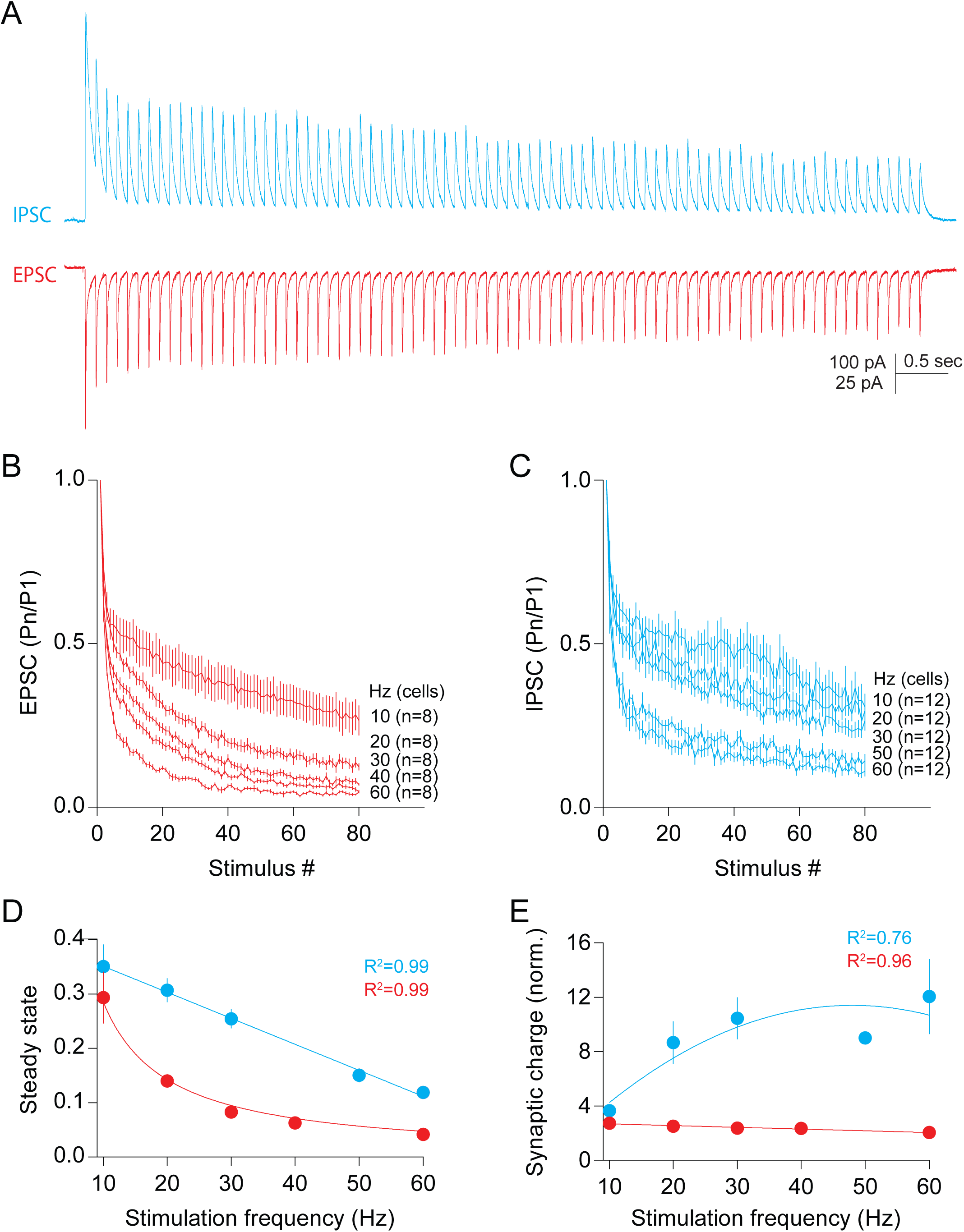
Differing dependence of short-term depression on stimulation frequency in excitatory and inhibitory synapses. (**A**) Current traces averaged from 10 responses evoked with extracellular stimulation (80 pulses at 10 Hz) under voltage clamp at −45 mV, to record IPSCs, or at 10 mV, to record EPSCs. (**B**) Amplitudes (normalized to the first pulse in the train) of the EPSCs evoked by pulse trains of stimulation at varying frequencies. (**C**) Normalized amplitudes of IPSCs evoked at different stimulation frequencies. (**D**) Average of normalized amplitudes at steady state (P60-P80) of EPSCs (red) and IPSCs (blue) evoked at different frequencies. (**E**) Synaptic charge of excitation (red) and inhibition (blue) recorded at steady state (P60-P80). Values were normalized to the product of synaptic charge of the first pulse in the train x the time duration. Fitting equation: 1/f (EPSCs) and line (IPSCs) in D; Line (EPSCs) and parabola (IPSCs) in E. Data from each cell are the averages of 10 repetitions.

Under the 1/f rule the steady state depression of the synaptic responses is proportional to the stimulation frequency. As a consequence, the total synaptic charge during a given time interval is independent of the frequency (Abbott et al., 1997; Tsodyks and Markram, 1997). As shown in figure 1E, the total synaptic charge (integrated between the 60^th^ and 80^th^ pulse and normalized to a single response) was constant over the 10 to 60 Hz range. In contrast, in the case of the compound IPSC, the relationship between stimulation frequency and the total synaptic charge per time interval was better fitted by a parabolic function (Parabola: R^2^ = 0.76, line: R^2^ = 0.62, regression; Probability that parabola is correct: 75.37%, Akaike’s Information Criteria; Figure 1E).

The lower level of sustained depression in inhibitory synapses suggests that the recovery from depression after sustained stimulation is faster at inhibitory synapses than at excitatory synapses. To test this prediction we evaluated the time taking to reach a new steady state when the stimulation frequency switches from 50 Hz to 10 Hz. The IPSC amplitude reached the new steady state much faster, with a shorter time constant (τ), than the EPSC amplitude (τ for IPSC: 0.27 ± 0.08 sec, n = 12; τ for EPSCs: 0.64 ± 0.09, n = 11; *U* = 16, p = 0.0013, M-W test; Figure S1). Altogether, these results confirmed that the depression-frequency relationship is different in excitatory and inhibitory synapses in layer 2/3 pyramidal cells of the visual cortex. The recovery from depression appears to be constant in the case of EPSC, and frequency-dependent in the case of IPSC.

### Use-dependent recovery from depression in inhibitory synapses from PV-INs

Extracellularly-evoked IPSCs primarily reflect the recruitment of PV-IN synapses. This encouraged us to investigate further with dual-patch whole-cell recordings from connected PV→Pyr pairs from G42 mice, in which GFP is expressed in a subpopulation of PV-INs. In these connected pairs we evoked a train of 30 action potentials in the PV-INs at a range of frequencies (from 10-60 Hz) and recorded the corresponding unitary IPSCs (uIPSCs) in the pyramidal cells. As shown in figure 2A, during the stimulation train the uIPSC amplitude rapidly depressed to reach a steady-state level by the 10^th^ pulse. As in the case of the evoked IPSCs (eIPSCs), the relationship between the average steady-state amplitude of the uIPSCs (measured as the last 10 responses) and the stimulation frequency did not follow the 1/f rule but was rather fitted by a linear function (Line: R^2^ = 0.97, 1/f: R^2^ = −4.01, regression; F(1,2) = 284.1, p = 0.0035, Extra sum-of-squares F test; Figure 2C). In a similar fashion, the relationship between synaptic charge per time interval of the uIPSCs and the stimulation frequency was not a constant, but was well fitted by a parabola (R^2^ = 0.98, regression; Figure 2D). Thus, like the eIPSCs, the depression of the uIPSCs does not follow the 1/f rule suggesting that the recovery rate of depleted synaptic resources depends on their use.

**Figure 2.**
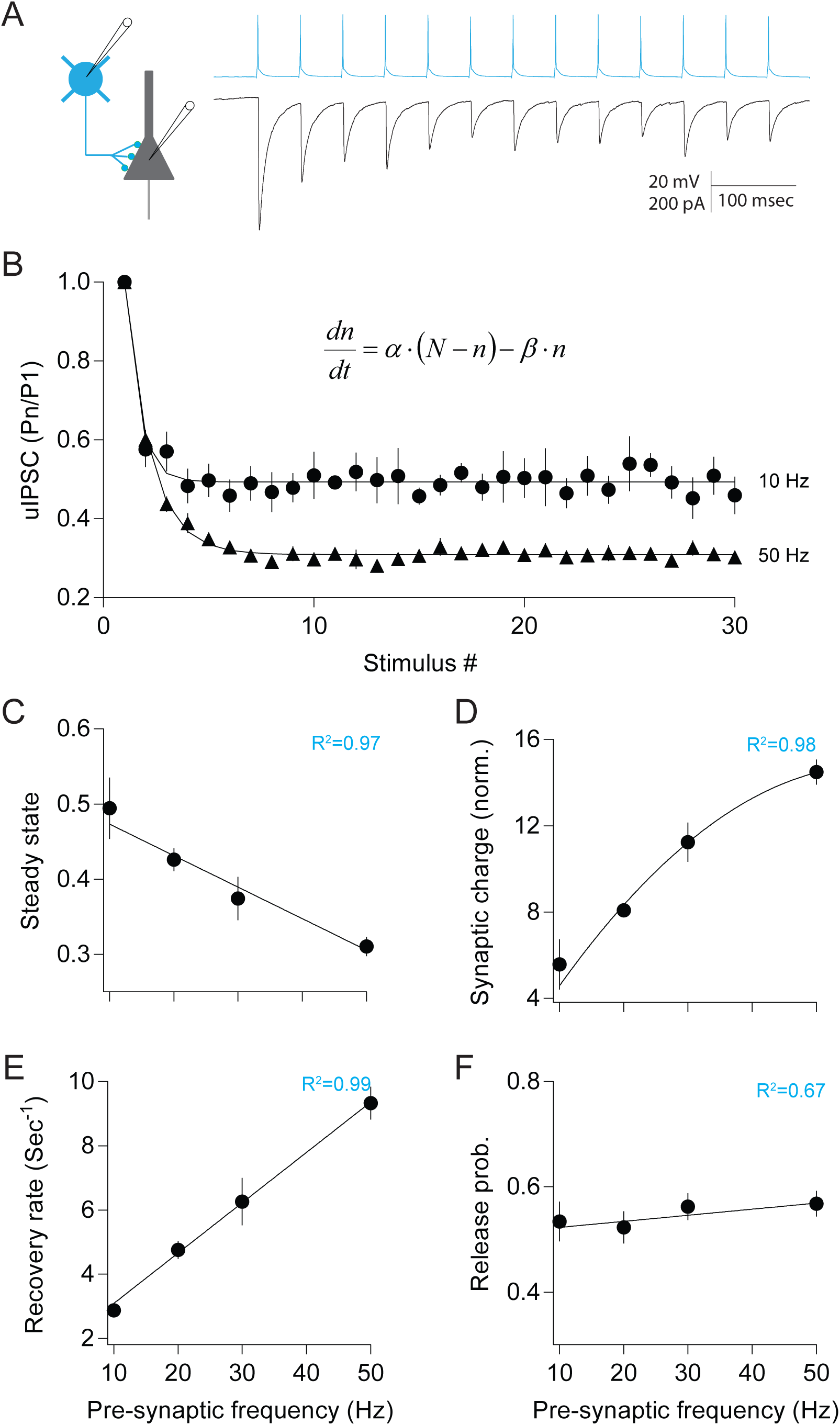
Dynamic synaptic replenishment in synapses made by parvalbumin interneurons (PV-INs) onto pyramidal cells (Pyr). (**A**) Recording from PV→Pyr pairs. Representative traces of action potentials (blue) and unitary inhibitory postsynaptic currents (grey) in a connected PV→Pyr pair in layer 2/3 of visual cortex in a G42 mouse. Action potentials were induced by current injection in the PV interneuron at 20 Hz. (**B**) Normalized amplitudes of uIPSCs evoked by 10 and 50 Hz of presynaptic stimulation. Curves were fitted by the function inset in the graph.(**C**) At the steady state (P20-P30) the amplitude of the uIPSCs were linearly related to the stimulation frequencies. (**D**) The average synaptic charge at the steady state increased parabolically with the stimulation frequency. (**E**) Rates of recovery from depletion, obtained by applying equation (1) to the data, increased linearly with presynaptic frequency. (**F**) Initial release probabilities were not affected by changing the presynaptic frequency. Fitting equation:line in C, E and F; Parabola in D. Sample size (cells): 7 (10Hz), 19 (20Hz), 10 (30Hz), 21 (50Hz). Data from each cell are the averages of 10-20 repetitions.

Next we evaluated the rate of recovery at different stimulation frequencies by fitting the depression of the uIPSCs during the stimulation train to equation (1) that describes a model in which synaptic resources are depleted and replenished (Wesseling and Lo, 2002).

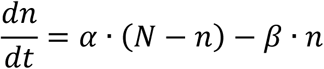

In this model, β is the intrinsic (initial) release probability, α is the rate of replenishment of synaptic resources (vesicles), n represents the pool of the available resources, and N is the maximal capacity of the resource pool. The results, plotted in figure 2E, indicate that at the steady state the rate of replenishment increased linearly with the stimulation frequency (R^2^ = 0.99, p = 0.0011, correlation). In contrast, the initial release probability remained constant (R^2^ = 0.67, p = 0.1792, correlation; Figure 2F). These results indicated that synaptic activity dynamically regulates the replenishment of synaptic resources.

To further confirm the dynamic regulation of the recovery from synaptic depression, we directly measured the recovery of uIPSCs following a conditioning stimulus. Depression was induced by conditioning with long stimulation train of 50 pulses delivered at either 20Hz or 50Hz; recovery was evaluated by single test pulses delivered at varied delay intervals (Δt). As shown in figure 3 the recovery trajectory as measured by the amplitude of the test uIPSCs was well fitted by an exponential curve at both cases 20 Hz and 50 Hz, and was faster in the case of conditioning with the 50 Hz stimulus (τ of 50 Hz: 187 ± 15 msec, n = 11; τ of 20 Hz: 412 ± 87 msec, n= 13; *U* = 30, p = 0.0154, M-W test; Figure 3B-C). These results confirmed that recovery is dynamically regulated by synaptic activity and that it is faster in more active synapses.

**Figure 3.**
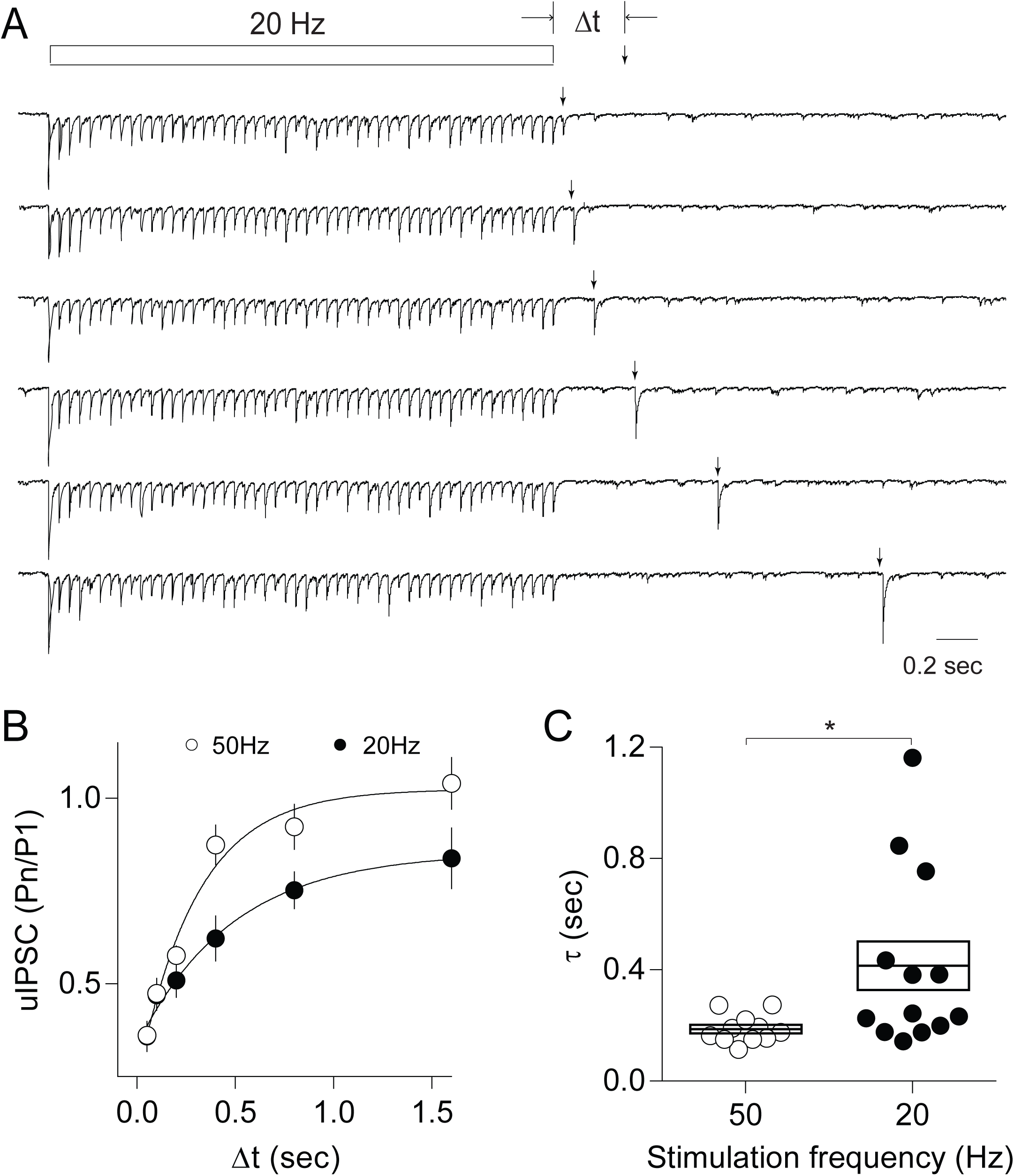
Synaptic recovery of PV→Pyr transmission is faster at a higher stimulation frequency.(**A**) Representative example of uIPSCs evoked by repetitive presynaptic firing at 20 Hz followed by a single presynaptic action potential evoked at intervals ranging from 50 msec to 1.6 sec. (**B**) Faster synaptic recovery after conditioning with 50 Hz (open symbols, n = 11) versus 20 Hz (filled symbols, n = 13) stimulation. The relative magnitudes of the uIPSCs were fitted by a single exponential function to obtain τ. (**C**) On average, the τ of recovery time after 50 Hz conditioning is significantly shorter than that after 20 Hz conditioning. *p = 0.0154, M-W test. Averages are represented by the horizontal middle lines in the boxes. Boxes indicate mean ± SEM. Data from each cell are the averages of 10 repetitions.

Previous studies have demonstrated developmentally-dependent postnatal changes in several important aspects of GABAergic transmission, including the number of synaptic contacts and release sites as well as the probability of release (Huang et al., 1999; Jiang et al., 2010; Morales et al., 2002). To determine whether activity-dependent dynamic recovery of PV→Pyr synapses also undergoes a maturational process, we evaluated the recovery rate and release probability from uIPSCs stimulated with 50Hz trains (50 pulses) and recorded from PV→Pyr pairs in both 2-week-old and 5-week-old mice (Figure S2A). The release probability, but not the recovery rate, was significantly reduced in the pairs from 5-week old mice (Release probability, 2W: 0.60 ± 0.03, n = 21; 5W: 0.46 ± 0.03, n = 13; t(32) = 2.942, p = 0.006, t-test; Recovery rate, 2W: 10.06 ± 0.65, n = 21; 5W: 9.53 ± 0.64, n =13; t(32) = 0.5530, p = 0.5841, t-test; Figure S2B-C). These results suggest that, unlike the release probability, the recovery rates remain constant during postnatal development.

### Rate coding of dynamic inhibitory recovery

Constant recovery and dynamic recovery of synaptic inhibition each will have distinct consequences for the firing of action potentials in pyramidal cells. In the case of constant recovery, increasing the stimulation rate of inhibitory inputs will barely increase the total synaptic charge per time unit and thus will barely affect firing rates. In the case of dynamic recovery, increasing the stimulation rate will allow an increase in inhibitory synaptic charge per time unit leading to reduced firing rates. We first explored these ideas in an all-active biophysical neuronal model from the Allen Cell Types Database (Model ID: 497232641) based on the morphology and electrophysiology of a pyramidal cell in layer 2/3 visual cortex (Cell ID: 477127614). The neuron model was driven by 3 independent excitatory inputs and 3 independent inhibitory inputs on the soma (Figure 4A) (see Methods for details on the assumptions of the response shape and release parameters). The postsynaptic conductance of each individual input was set to 4 nS in the case of inhibition (based on unpublished data) and to 5 nS in the case of excitation. These inputs were activated with Poisson trains of different frequencies (Figure 4B). Ten repetitions per frequency with new Poisson trains in each trial were performed, and firing rates were measured from whole trials. In the absence of inhibition, increasing the frequency of the excitatory inputs increased the firing rates to a limit of about 60 Hz (Figure 4B). The recruitment of inhibitory inputs with constant recovery reduced the firing only slightly, and increases in the inhibitory frequency had a modest effect on the model cell firing rates (Figure 4C-E). In contrast, the recruitment of inhibitory inputs with dynamic recovery had a larger impact on firing, and increasing the inhibitory frequency further reduced model cell firing in a multiplicative manner (Figure 4C-E).

**Figure 4.**
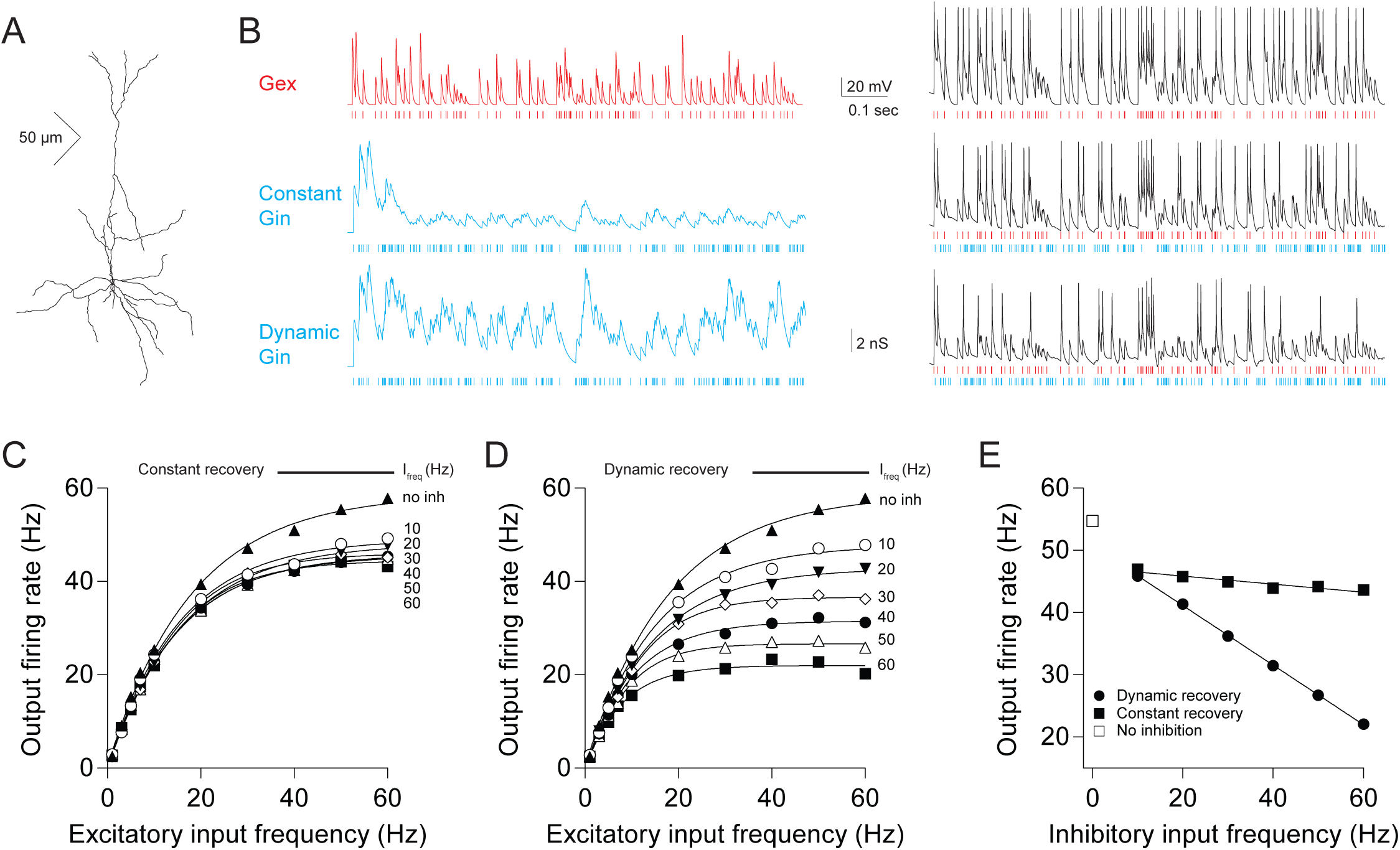
Dynamic recovery of inhibitory transmission enables linear gain control of firing in a neuronal model. (**A**) The 3-dimenstion structure of the all-active neuron model (Model ID: 497232641 at the Allen Cell Types Database) used in the simulation. (**B**) Simulation of excitatory and inhibitory inputs with Poisson trains. Left: examples of excitatory synaptic conductance (Gex, 30 Hz) and inhibitory synaptic conductance (60 Hz) with either constant rate of recovery (constant Gin) or dynamic rate of recovery (Dynamic Gin). Right: example traces of membrane potential during activation of excitatory inputs only (top), activation of excitatory and inhibitory synapses with constant recovery rate (middle), or activation of excitatory synapses and inhibitory synapses with dynamic recovery (bottom). The vertical lines at the bottom of traces for conductance and membrane potential indicate the activation timing of excitatory synapses (red) and inhibitory synapses (blue). (**C**) Modest effects of inhibition with constant-recovery on firing rates of the model pyramidal cell. Plotted are the average firing rates elicited by different frequencies of the excitatory input, and in the absence of (filled up-pointing triangles) or in conjunction with (other symbols) stimulation of inhibitory inputs at the indicated frequencies. (**D**) Inhibition with dynamic synaptic recovery resulted in greater attenuation of the firing rates in the model cell. Excitatory input vs. firing output curves as in C. (**E**) Inhibitory inputs vs. firing output averaged at the 40-60 Hz excitatory activity (horizontal bar in C and D), indicating inhibition with dynamic recovery linearly reduced neural firing rate. Fitting equation: sigmoid in C and D; Line in E. Data points in C to E are the averages from 10 trials.

Next we confirmed the differential effects of these two types of inhibitory synapses in actual neurons using dynamic clamp to simulate excitatory and inhibitory currents. Currents trains based on excitatory conductance, as well as inhibitory conductance with either dynamic or constant recovery, were injected into layer 2/3 pyramidal cells and the effect on the membrane potential was recorded (Figure 5A). The excitatory and inhibitory conductance, representing 3 independent excitatory inputs and 3 independent inhibitory inputs, respectively, had the same response shape and release parameters as that in the neural model. The conductance of the excitatory input was set to 1.5 times the action potential threshold, and the inhibitory conductance was set such that inhibitory inputs (30 Hz) with constant recovery could reduce the firing induced by excitatory inputs (30 Hz) by about 30%. Stimulation trains of a given frequency were run once in each cell. The inter-train interval was set from 20-30 sec. In the absence of inhibition, increasing the excitation frequency resulted in increased firing that reach a plateau of 18.69 Hz, with a stimulus frequency over 20 Hz. We then tested the effects on firing elicited by excitation at 30 Hz. As shown in figure 5B-C, as in the case of the model neuron, inhibitory synapses with dynamic recovery imposed multiplicative inhibition on the firing activity of the pyramidal neuron, while inhibitory synapses with constant recovery had only a modest effect on firing rate which was largely independent of the inhibitory frequency (Dynamic recovery: R^2^ = 0.92, p = 0.0027; Constant recovery: R^2^ = 0.49, p = 0.1212; Correlation; Figure 5C). Thus, only inhibitory synapses endowed with dynamic recovery can support a frequency dependent graded attenuation of pyramidal cell firing.

**Figure 5.**
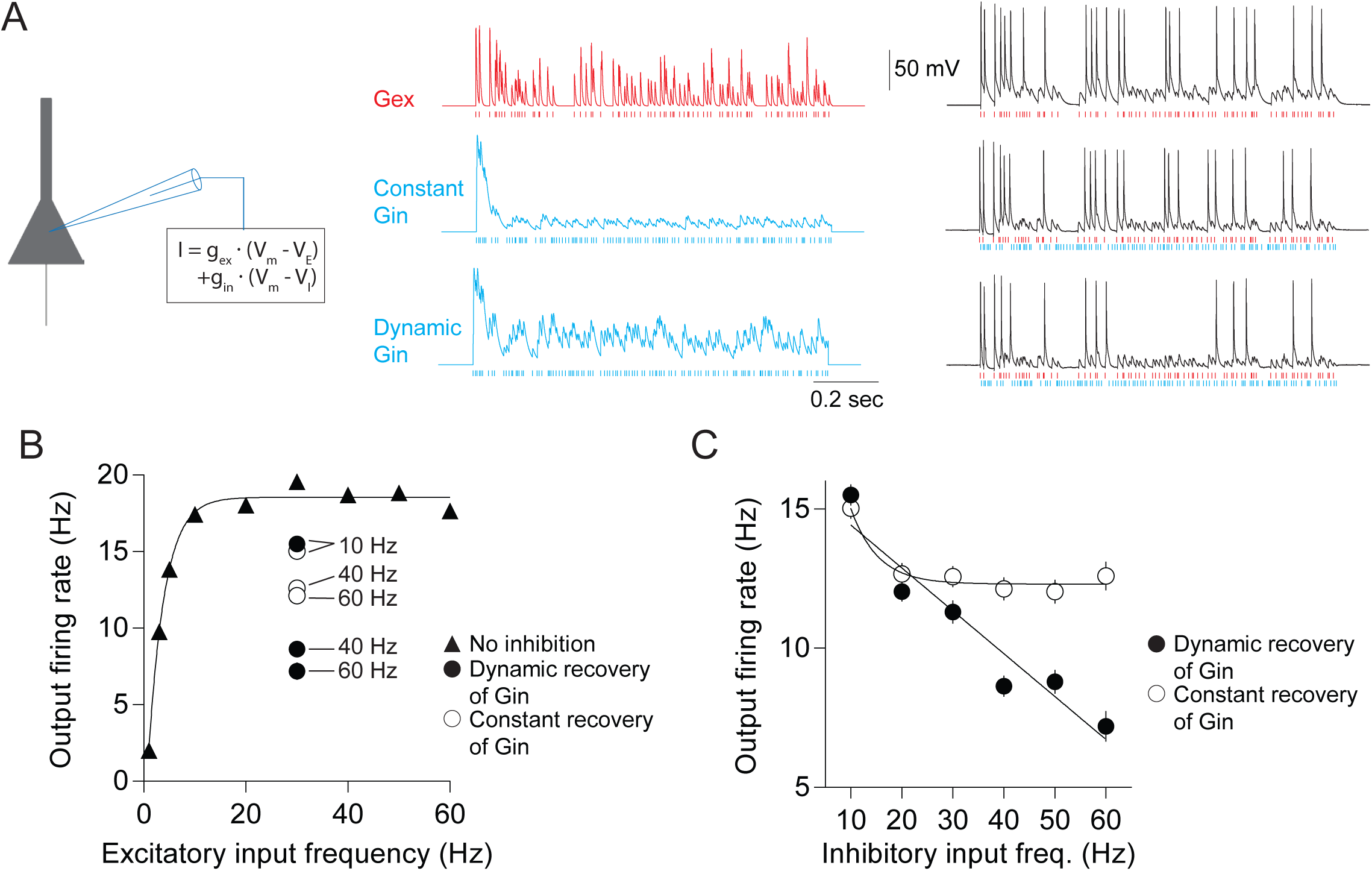
Dynamic vesicle recovery enabled linear gain control of inhibitory transmission in neurons with dynamic clamping. (**A**) Left: illustration of dynamic clamping in real neurons. Middle: excitatory conductance and inhibitory conductance with either constant recovery or with dynamic recovery were injected into neurons using dynamic clamp. Right: examples of membrane potentials evoked by excitatory inputs only, by excitatory inputs and inhibitory inputs with constant recovery, or by excitatory inputs and inhibitory inputs with dynamic recovery. The vertical lines at the bottom of traces for conductance and membrane potential indicated the activation timing of excitatory synapses (red) and inhibitory synapses (blue). (**B**) Excitatory input vs. firing output functions without inhibition, with dynamic-recovery inhibition (10, 40, and 60 Hz) and with constant-recovery inhibition (10, 40, and 60 Hz). (**C**) Relationship of inhibitory input vs. firing output, indicating inhibition with dynamic recovery linearly reduced neuronal firing. The frequency of excitation in C is 30Hz. Sample size: n = 30 cells. Fitting equation: sigmoid in B; Line (Dynamic recovery of Gin) and one phase decay (Constant recovery of Gin) in C.

Finally, we asked whether real inhibitory synapses have the capacity to reduce pyramidal cell firing in a frequency-dependent manner. To do this we recorded from pyramidal cells and combined dynamic clamp (to mimic trains of excitatory conductance) with optogenetic recruitment of inhibitory cells in slices prepared from mice expressing channelrhodopsin-2 in PV-INs (see Methods for details). To validate the optogenetic approach we first confirmed that these PV-INs can produce action potentials at frequencies as high as 50Hz when stimulated with pulses of blue light (Figure S3A). We also confirmed that upon repetitive stimulation the light-induced IPSCs recorded in pyramidal cells do behave like the unitary IPSCs recorded from PV→Pyr pairs. The light induced IPSCs exhibited depression (Figure S3B), with a steady state amplitude that decreases linearly with the light pulse frequency and a synaptic charge that increases in a parabolic manner (Steady state: R^2^ = 0.99, p = 0.0002, correlation; Synaptic charge: R^2^ = 0.99, regression; Figure S3C-D). Moreover, fitting the data with equation (1) yielded estimated recovery rates that depend linearly on light frequency (R^2^ = 0.99, p < 0.0001, correlation; Figure S3E), though release probability was also found to depend on frequency slightly (R^2^ = 0.93, p = 0.0087, correlation; Figure S3F).

To evaluate the impact of recruiting inhibition mediated by PV-INs, we first made the pyramidal cell fire by injecting trains of excitatory currents (30 Hz, the same excitatory conductance as that in figure 5C) in the dynamic-clamp mode while delivering a Poisson train of light pulses (Figure 6B), with the light intensity adjusted to induce IPSCs of approximately 1 nA when holding at 0 mV. As shown in figure 6C, increasing the average frequency of the light pulses increased the attenuation of the firing of the pyramidal cell. Moreover, the firing rate of the pyramidal cell was decreased linearly as a function of the frequency of the recruitment of PV-IN synapses (R^2^ = 0.95, p = 0.0052, correlation; Figure 6D). These results indicate that synapses made by PV-INs, endowed with dynamic recovery, can exert a graded control of pyramidal cell spiking based on the firing rate of the PV-INs.

**Figure 6.**
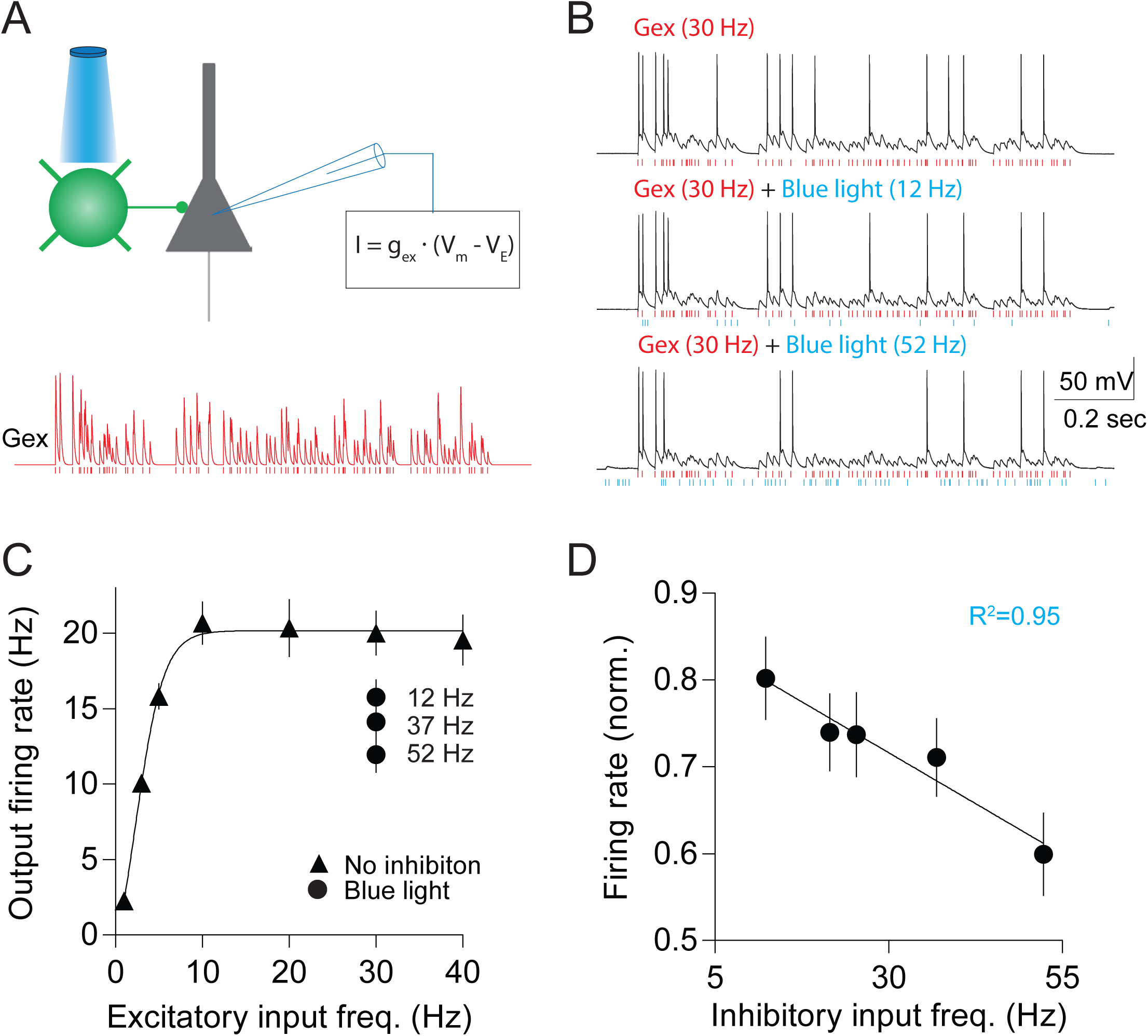
Optogenetically activated inhibition exhibited linear gain control. (**A**) Inhibitory inputs from light-controlled parvalbumin interneurons and excitatory inputs from dynamic clamping. Bottom trace is an example of the excitatory conductance while the vertical red lines indicate the activation timing of excitatory synapses. (**B**) Example of membrane potentials evoked by excitatory inputs only, excitatory inputs and inhibitory inputs at 12 Hz, or excitatory inputs and inhibitory inputs at 52 Hz. The vertical lines at the bottom of the membrane potential traces indicate the activation timing of excitatory synapses (red) and blue light (blue). (**C**) Excitatory input vs. firing output without or with inhibition (12, 37, and 52 Hz). (**D**) Averaged firing rate (normalized to the firing rate evoked by 30 Hz excitation only) vs. the frequency of inhibitory inputs, indicating inhibition from PV cells linearly reduced neural activity in pyramidal cells. The frequency of excitation in D is 30 Hz. Sample size: n = 15 cells. Fitting equation: sigmoid and line in B and C, respectively.

## DISCUSSION

In this study we revealed that the recovery from short-term depression in inhibitory synapses made by PV-INs onto pyramidal cells is activity-dependent, such that increasing the frequency of presynaptic activity also increases recovery rates. As a result, the inhibitory output, measured as the total charge per unit time, varies linearly with the presynaptic frequency. This dynamic synaptic recovery allows these inhibitory synapses to multiplicatively reduce pyramidal cell firing rates over a wide range of frequencies.

The exact mechanisms underlying such dynamic recovery at inhibitory synapses made by PV interneurons remain unknown. One attractive possibility is the accumulation of intracellular Ca^2+^. In many synapses the replenishment of the readily releasable pool of vesicles (RRP) can be accelerated by activity-induced calcium accumulation (Dittman and Regehr, 1998; Fuhrmann et al., 2004; Gomis et al., 1999; Hosoi et al., 2007; Sakaba and Neher, 2001; Smith et al., 1998; Stevens and Wesseling, 1998; Wang and Kaczmarek, 1998). Repetitive stimulation causes calcium to accumulate in presynaptic terminals (Kreitzer et al., 2000), potentially at levels commensurate with the stimulation frequency. It must be noted that parvalbumin, being a slow calcium buffer, may contribute to the activity-dependent accumulation of Ca^2+^ at presynaptic terminals. However, in PV knockout mice the IPSCs evoked by repetitive stimulation are altered only in the early part of the train, not in the steady state, when the rate of replenishment becomes a limiting factor of the IPSC amplitude (Caillard et al., 2000; Eggermann and Jonas, 2011; Orduz et al., 2013; Vreugdenhil et al., 2003).

The genetic ablation of downstream Ca^2+^-sensing molecules can impair the activity-dependent recovery of the vesicle pool in various synapses. These molecules include the Ca^2+^- Calmodulin-Munc13-1 complex found at the calyx of Held synapses (Lipstein et al., 2013; Sakaba and Neher, 2001) as well as synaptotagmin 2 and 7 (Syt-2 and Syt-7), which are found at basket cell to Purkinje cell synapses (Chen et al., 2017a; Chen et al., 2017b; Turecek et al., 2017) and in cultured hippocampal synapses (Liu et al., 2014), but not at the calyx of Held synapses (Luo and Sudhof, 2017). Of these calcium-sensor molecules, a particularly attractive candidate to mediate dynamic recovery in the cortex is Syt-2, which is highly and selectively expressed in the terminals of cortical PV-INs (Sommeijer and Levelt, 2012).

Concerning the functional consequences of dynamic recovery, a widely accepted role of the network of PV-INs in sensory cortices is to control the gain of pyramidal cell responses to sensory stimulation (Atallah et al., 2012; Hofer et al., 2011; Packer and Yuste, 2011). According to this view, the high interconnectivity between PV-INs and pyramidal cells (Gu et al., 2013; Holmgren et al., 2003; Packer and Yuste, 2011) allows PV-INs to sense the activity of a population of pyramidal cells and to broadcast back a proportionate and graded inhibitory signal to multiple pyramidal cells. Relevant to this function is our finding that increasing PV-IN firing can reduce PC firing in a linear fashion (Figure 6D), which in a model can only be accomplished by inhibitory synapses. Thus, dynamic recovery complements other unique functional specializations of PV-INs, including the expression of Kv3 channels in the soma that allows a large dynamic range of action potential firing (with rates >200Hz) and in the axon terminals, to allow high rates of release (Goldberg et al., 2005). In addition, PV-INs also contribute to the generation of gamma-oscillations (Bartos et al., 2007; Sohal et al., 2009), and it is conceivable that dynamic recovery at PV synapses could be essential for maintaining these high frequency oscillations. Besides, dynamic recovery at inhibitory synapses resulted in relativity less depression at high frequencies of activity, which tilted the balance of excitation and inhibition towards inhibition. This shift of E/I balance may stabilize network activity (Varela et al., 1999). Since imbalanced excitation and inhibition and altered gamma-oscillation have been linked to autism, schizophrenia, and many other neurodevelopmental disorders (Hussman, 2001; Lisman, 2012; Rojas and Wilson, 2014; Rubenstein and Merzenich, 2003; Simon and Wallace, 2016), this work may provide important insight into the etiology of these conditions.

## EXPERIMENTAL PROCEDURES

### Animals

2-5 week old C57BL/6 mice, 2-5 week old G42 mice, and 3-8 week old Parvalbumin (PV)-Cre mice (Stock #: 008069, Jackson laboratory) of either sex reared in normal light/dark 12-hour cycles were used in these studies. Mice used at the Hussman Institute for Autism were housed and cared for by the AAALAC accredited program of the University of Maryland School of Medicine. All procedures were approved by the Institutional Animal Care and Use Committee (IACUC) at the Johns Hopkins University. Procedures performed at the Hussman Institute for Autism were reviewed and approved by the IACUCs at the University of Maryland School of Medicine and the Hussman Institute for Autism.

### Channelrhodopsin-2(ChR2) Viral injection

At age 3-4 weeks, PV-Cre mice were anesthetized with 1-3% isoflurane mixed with O^2^ and transcranially injected bilaterally into layer 2/3 of the visual cortex (coordinates: Bregma 3.6, lateral 2.5, depth 0.36) with 1.5 μl adeno-associated virus containing ChR2 and yellow fluorescence protein or mCherry as a marker (Addgene26973 or Addgene20297, Penn Vector Core). Mice were allowed to recover on a heated surface and were returned to the animal colony, where they remained for 2-4 weeks to allow for optimal ChR2 expression before experimental procedures were initiated.

### Preparation of visual cortical slices

Visual cortical slices (300 μm) were cut as described previously (Bridi et al., 2017; Gu et al., 2013; Huang et al., 2013; Huang et al., 2012) in ice-cold dissection buffer containing the following (in mM): 212.7 sucrose, 5 KCl, 1.25 NaH_2_PO_4_, 10 MgCl_2_, 0.5 CaCl_2_, 26 NaHCO_3_, 10 dextrose, bubbled with 95% O_2_/5% CO_2_, pH 7.4. Slices were transferred to normal artificial CSF (ACSF) at 30 °C for 30 min. Slices were recovered at room temperature for at least an additional 30 min before recording. Normal ACSF was similar to the dissection buffer except that sucrose was replaced by 124 mM NaCl, MgCl_2_ was lowered to 1 mM, and CaCl_2_ was raised to 2 mM. All recordings were performed at 30°C.

### Visualized whole-cell recording

Glass pipettes (4-6 MΩ) filled with one of three types of internal solution were used, depending on the experiment: a K-based internal solution (in mM: 130 KGluconate, 10 KCl, 0.2 EGTA, 10 HEPES, 0.5 NaGTP,4 MgATP, and 10 Na_2_-Phosphocreatine; pH adjusted to 7.25 with KOH, 280–290 mOsm), a CsCl-based internal solution (in mM: 120 CsCl, 8 NaCl, 10 HEPES, 2 EGTA, 5 QX-314, 0.5 Na_2_GTP, 4 MgATP, and 10 Na2-Phosphocreatine; pH adjusted to 7.25 with CsOH, 280–290 mOsm), and a CsGluconate-based internal solution(in mM: 130 CsGluconate, 8 KCl, 10 EGTA, 10 HEPES and 10 QX-314; pH adjusted to 7.25 with CsOH, 280–290 mOsm). Patch clamping properties were monitored with 100-msec negative voltage or current commands (−2 mV and −40 pA for voltage and current clamp, respectively) delivered every 20-30 seconds. Cells were excluded if series resistance changed > 15% over the experiment. Only cells with series resistance < 20 MΩ, and input resistance > 100 MΩ were included. Data were filtered at 2 kHz and digitized at 10 kHz using Igor Pro (WaveMetrics, Portland, OR).

For recording excitatory and inhibitory postsynaptic currents (EPSCs and IPSCs) in pyramidal cells of layer 2/3, with extracellular stimulation at layer 4, whole-cell voltage clamping was done with the CsGluconate-based internal solution. To isolate EPSCs and IPSCs, membrane potentials were held at the reversal potentials of EPSCs and IPSCs (−45 mV and +10 mV, respectively). In some cases, only IPSCs were recorded with the CsCl-based internal solution, at a holding potential of −60mV, and in the presence of APV (100 μM) and CNQX (20 μM) in the ACSF to block excitatory transmission. Extracellular stimulation was delivered through one concentric bipolar electrode (125 μm diameter) (FHC, Bowdoin, ME) placed in layer 4 of visual cortex. Stimulus intensity was adjusted to evoked simple-waveform postsynaptic currents.

For recording unitary IPSCs (uIPSCs) in pyramidal cells, dual whole-cell recordings were made from pairs of GFP-expressing PV interneurons and pyramidal neurons. Glass pipettes filled with K-based internal solution were used to record from presynaptic PV interneurons. Pipettes filled with CsCl-based internal solution were used to record from postsynaptic pyramidal neurons. uIPSCs were recorded in pyramidal neurons in voltage clamp mode at −60 mV, and were evoked by suprathreshold somatic square-current injection (2 msec) in presynaptic fast-spiking PV interneurons (Gao et al., 2017; Gu et al., 2013; Jiang et al., 2010).

For recording action potentials in PV interneurons expressing ChR2, whole-cell current clamp recordings were made in PV interneurons using the K-based internal solution. Blue light at 455 nm or 470 nm with a duration of 2 msec was generated by collimated LED light sources (Thorlabs, Newton, New Jersey) controlled by an LED driver (Thorlabs, Newton, New Jersey).

For recording ChR2-evoked IPSCs in pyramidal cells, whole-cell voltage clamp recordings were made in pyramidal cells of layer 2/3 in visual cortical slices, in which ChR2 was expressed in PV-Cre interneurons. The K-based internal solution was used, and membrane potentials were held at 0 mV.

For dynamic clamp experiments, whole-cell current clamp recordings were made from layer 2/3 pyramidal cells with the K-based internal solution. Dynamic clamp was performed with modified StdpC software using source code provided by Dr. Thomas Nowotny (University of Sussex) (Kemenes et al., 2011), in which excitatory and inhibitory conductance were pre-generated. The excitatory and inhibitory conductance, composed of 3 independent excitatory inputs and 3 independent inhibitory inputs, respectively, had the same response shape and release parameters as that in the simulation of the neural model. The amplitude of conductance was determined on a cell-by-cell basis using the following rules: The conductance of the excitatory input was set to 1.5 times the action potential threshold, and the inhibitory conductance was set such that inhibitory inputs (at 30 Hz) with constant recovery could reduce firing induced by excitatory inputs (at 30 Hz) by about 30%. Stimulation trains were run once in each cell. The duration of each train was 1 sec. The inter-train interval was set from 20-30 sec. Firing rates were measured from whole trials.

### Data analysis

Vesicle release probability and recovery rate were obtained following the method developed by Wesseling and Lo (2002). Briefly, the amplitudes of uIPSCs were fitted by two functions (2 and 3) derived from the equation (1) below:

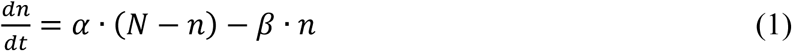

Where, *α* denotes the recovery rate; *β* denotes the release probability; *n* denotes vesicles available for release and *N* denotes total vesicles in ready release pool (Wesseling and Lo, 2002).

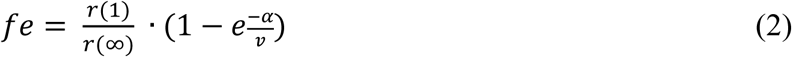

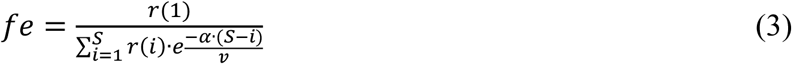

Where, *fe* is the initial release probability; *r*(1) is the amount of transmitter released by the first stimulus of the train; *r*(∞) is the release at the steady-state of the train; *v* is the stimulation frequency of the train; *S* is the total number of the stimuli in the train; *i* is the *i*th stimuli in the train.

The total synaptic charges during steady state of the train were measured by integrating postsynaptic currents in the last 10 or 20 stimuli of the steady state and were then normalized by the product of the synaptic charge of the first pulse in the stimulation train and the time duration of the stimuli. The curve of normalized synaptic charges was fitted by a parabolic function of presynaptic firing frequency (*v*):

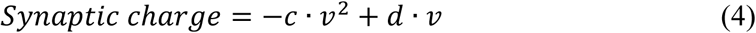

Firing rates in the model neuron and real neurons were calculated by measuring the number of action potentials in the whole trace and divided by the time duration.

### Simulation

Simulation was ran in Python (https://www.python.org) with the Neuron simulator (http://www.neuron.yale.edu). An all-active realistic biophysical neuronal model (Model ID:497232641, Allen Cell Types Database (2017)) was connected by 3 excitatory synapses and 3 inhibitory synapses at the soma (Gouwens et al., 2018). All synapses were activated by independent Poisson-distributed stimulation. Excitatory and inhibitory synaptic conductances (*G*) were described by rising time (τ1) and decay time (τ2) as below:

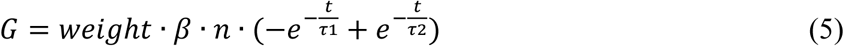

For excitatory synapses, τ1 and τ2 were set as 0.3 msec and 3 msec, respectively (Gao et al., 2017; Hausser and Roth, 1997). For inhibitory synapses, τ1 and τ2 were set as 0.6 msec and 15 msec, respectively (based on unpublished data). Vesicle release and recovery mechanisms as defined by the equation (1) were included in both excitatory and inhibitory synapses with parameters obtained from experimental data. The total vesicles in the ready release pool, *N*, was set as 1. The release probability (*β*) and recovery rate (*α*) for excitatory synapses were 0.56 and 9 sec^-1^, respectively. The release probability and recovery rate for inhibitory synapses with constant recovery were 0.56 and 3.099 sec^-1^, respectively. In the inhibitory synapses with dynamic vesicle recovery, the recovery rate (*α*) was changed as a function of presynaptic firing frequency (*v*):

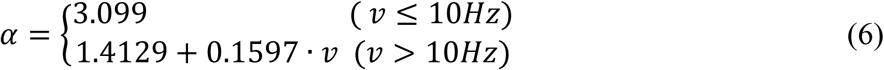

Synaptic currents (*I*) were described by the following function:

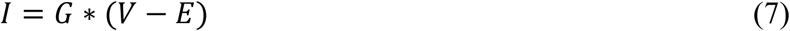

Where, *V* is the membrane potential; *E* is the reversal potential which was set as 0 mV and −70 mV for excitatory and inhibitory synapses, respectively.

### Statistical analysis

Graphpad Prism 6 software (Graphpad Prism) was used to perform all statistical analyses. For two-group comparisons, significance was examined by unpaired two-tailed t-tests or Mann-Whitney (M-W) tests based on the normality of data set using the D’Agostino-Pearson omnibus normality test. Nonlinear regressions were performed to fit the data. Pearson correlation or nonparametric Spearman correlation were performed to assess the correlation between two variables. The Extra sum-of-squares F test or Akaike’s Information Criteria (ALCc) was used to compare which of two models fits best. Significance was considered to be p < 0.05. All data are presented as mean ± SEM.

## ACKNOWLEDGMENTS

We thank Dr. H.-K. Lee for valuable discussions and comments, and Dr. J. L. Whitt for help in viral injection. This work was supported by NIH grants R01EY012124 to A.K. and Hussman Foundation grants HIAS18001 to S.H.

## AUTHOR CONTRIBUTIONS

Conceptualization, S.H. and A.K.; Methodology, S.H. and A.K.; Software, S.H., Formal Analysis, S.H., M.S.B. and A.K.; Investigation, S.H., M.S.B.; Writing – Original Draft, S.H. and A.K.; Writing – Review & Editing, S.H., M.S.B. and A.K.; Visualization, S.H. and A.K.; Funding Acquisition, S.H. and A.K.; Supervision, S.H. and A.K.

---

**Figure S1.**
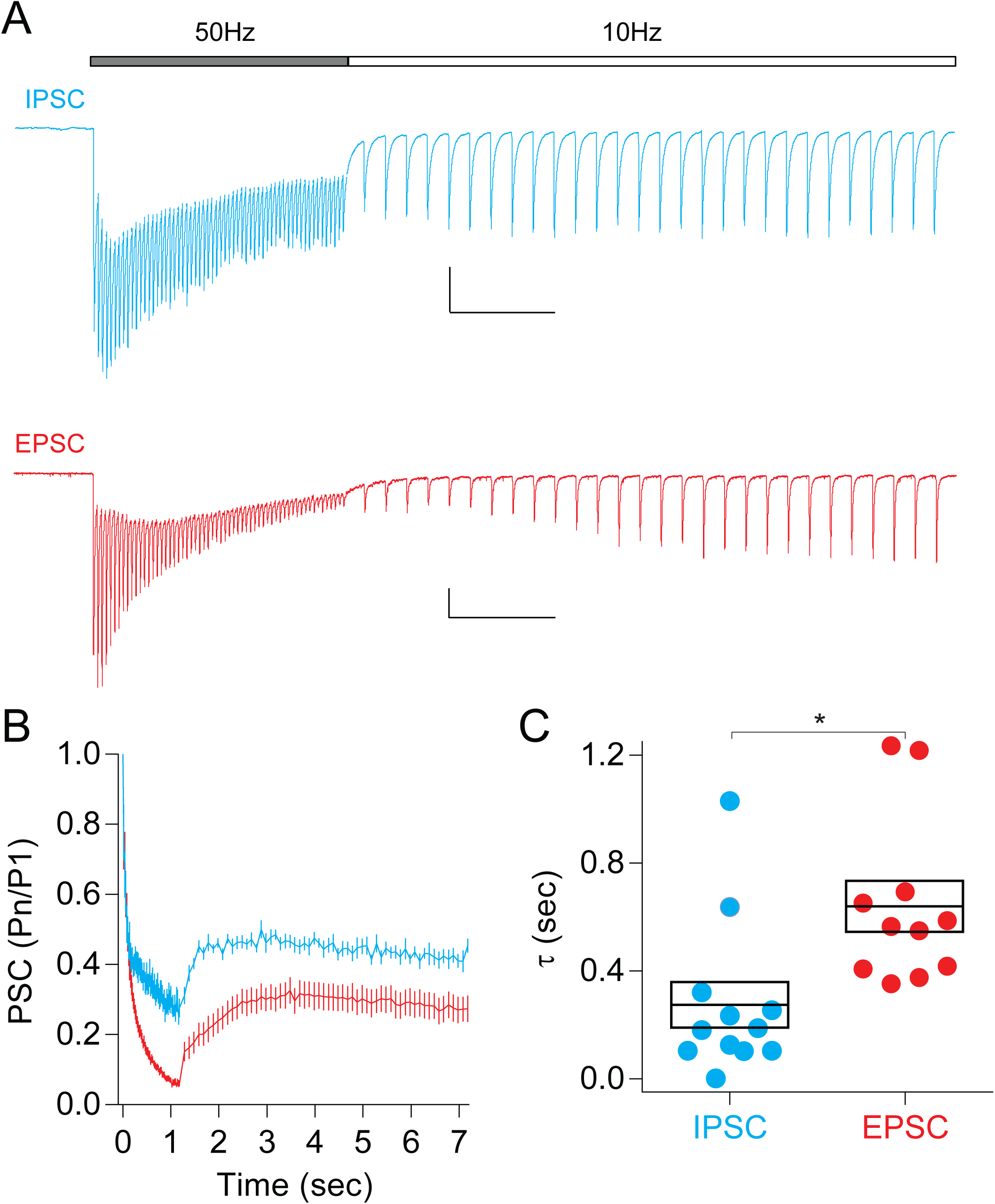
Related to Figure 1. Transition from high to low frequency stimulation in excitatory and inhibitory synapses. (**A**) Representative traces of IPSCs and EPSCs recorded while stimulation frequency switched from 50 Hz to 10Hz. (**B**) Normalized amplitudes of IPSCs (blue) and EPSCs (red). (**C**) Time constant τ during transition from 50 Hz to 10 Hz in IPSCs was smaller than that in EPSCs. *p = 0.0013, M-W test. Scale bars in A; 200pA (top), 50 pA (bottom) and 0.5 sec. In these experiments the IPSCs were recorded with CsCl based internal solution. Cells held at −60 mV. Sample size (cells): n = 12 in IPSCs and n = 11 in EPSCs. Data from each cell are the averages of 10 repetitions.

**Figure S2.**
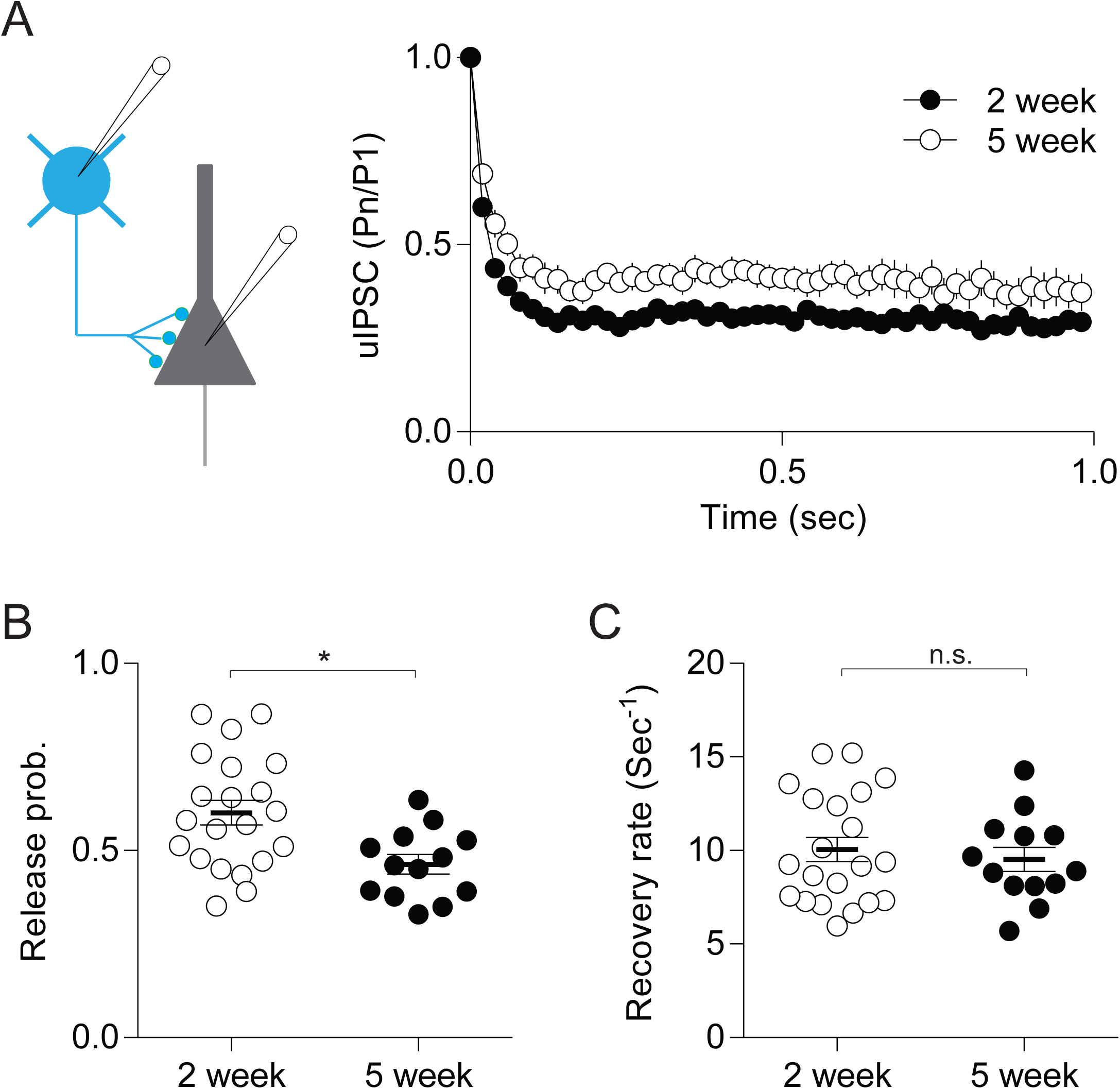
Related to Figure2. Reduced vesicle release but not recovery in inhibitory synapses from PV interneurons to pyramidal cells during postnatal development. (**A**) Normalized uIPSCs evoked by 50 Hz presynaptic stimulation in 2-week-old and 5-week-old G42 mice. (**B**) Release probability in inhibitory synapses during postnatal development. *p = 0.006, t-test. (**C**) Recovery rate in inhibitory synapses during postnatal development. n.s.: p = 0.5841, t-test. Sample size(cells): n = 21 (2 week) and n = 13 (5 week). Data from each cell are the averages of 20-30 repetitions.

**Figure S3.**
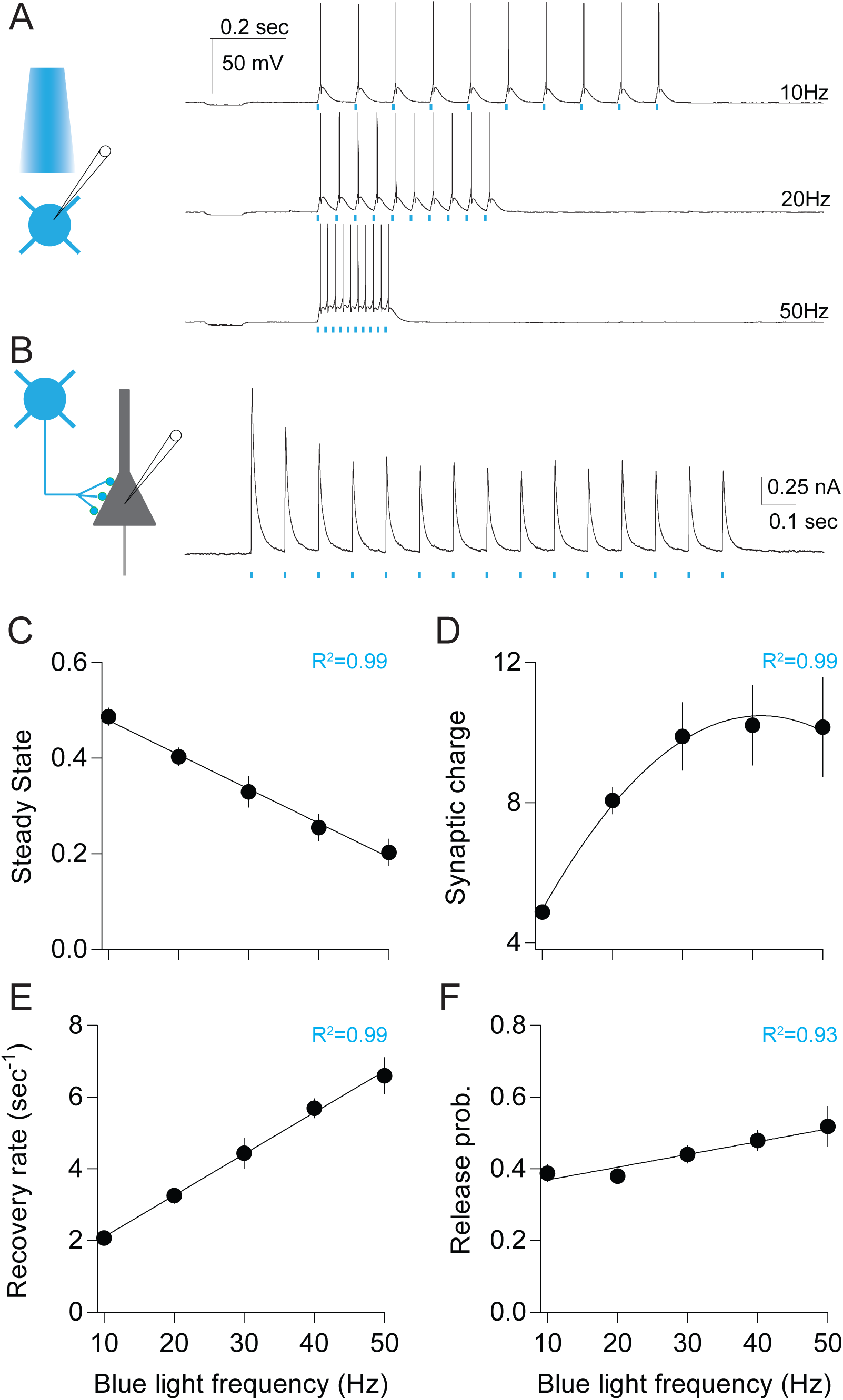
Related to Figure 6. Dynamic vesicle replenishment in optogentically activated inhibitory synapses from PV interneurons to pyramidal cells. (**A**) Action potentials could be induced at frequencies as high as 50 Hz in PV-INs expressing ChR2. Blue bars indicated timing of blue light stimulation. (**B**) Example trace of IPSCs in a pyramidal cell evoked by blue light at 10 Hz. (**C**) Normalized IPSCs at steady states with different frequencies of light stimulation. (**D**) Normalized steady-state synaptic charges. (**E**) Vesicle recovery rate in inhibitory synapses activated by optogenetic stimulation. (**F**) Vesicle release probability in inhibitory synapses activated by optogenetic stimulation. Sample size: n = 11 cells. Fitting equation: line in C, E and F; Parabola in D. Data from each cell are the averages of 6-7 repetitions.

